# High-resolution disease phenotyping reveals distinct resistance strategies of wild tomato crop wild relatives against *Sclerotinia sclerotiorum*

**DOI:** 10.1101/2024.05.07.592883

**Authors:** Severin Einspanier, Christopher Tominello-Ramirez, Mario Hasler, Adelin Barbacci, Sylvain Raffaele, Remco Stam

## Abstract

Besides the well-understood qualitative disease resistance, plants possess a more complex quantitative form of resistance: quantitative disease resistance (QDR). QDR is commonly defined as a partial but more durable form of resistance and, therefore, might display a valuable target for resistance breeding. The characterization of QDR phenotypes, especially of wild crop relatives, displays a major bottleneck in deciphering QDR’s genomic and regulatory background. Moreover, the relationship between QDR parameters, such as infection frequency, lag phase duration, and lesion growth rate, remains elusive. High hurdles for applying modern phenotyping technology, such as the low availability of phenotyping facilities or complex data analysis, further dampen progress in understanding QDR. Here, we applied a low-cost phenotyping system to measure lesion growth dynamics of wild tomato species (e.g., *S. pennellii* or *S. pimpinellifolium*). We provide insight into QDR diversity of wild populations and derive specific QDR strategies and their crosstalk. We show how temporally continuous observations are required to dissect end-point severity into functional resistance strategies. The results of our study show how QDR can be maintained by facilitating different defense strategies during host-parasite interaction and that the capacity of the QDR toolbox highly depends on the host’s genetic context. We anticipate that the present findings display a valuable resource for more targeted functional characterization of the processes involved in QDR. Moreover, we show how modest phenotyping technology can be leveraged to help answer highly relevant biological questions.

## 1. Introduction

### Quantitative disease resistance in plants

Plant resistance is commonly divided into two concepts with fundamental differences: qualitative and quantitative resistance [1,2]. While qualitative disease resistance provides a highly effective race-specific resistance, quantitative disease resistance (QDR) is a broad-range yet incomplete resistance [2,3]. Qualitative resistance is driven by major race-specific resistance genes (R-genes). They often lead to complete and easily observable resistance and were the dominant research focus for disease resistance breeding programs. However, reports of R-genes losing their efficacy against pathogens have increased recently, and major resistance genes have not been identified for many so-called necrotrophic plant pathogens, like *Botrytis cinerea* or *Sclerotinia sclerotiorum* [3–7]. Commonly, degrees of QDR can’t be divided into discrete classes. Quantitative resistance phenotypes are continuously distributed and can only be explained by highly integrated, polygenic regulatory mechanisms [8]. Moreover, QDR can manifest itself in several ways, ranging from differences in infection frequency on the leaf or delayed onset of infection to stalled lesion growth. Numerous studies documented wide distributions of QDR phenotypes against necrotrophic pathogens in both natural and domesticated plant populations, yet the relations of different QDR phenotypes have not yet been studied in detail [1–3,8–12]. Recent reports summarized the diversity in functional QDR, arguing that QDR might be influenced by many independent components such as regulation as a pleiotropic side-effect, weak R-genes, involvement in defense signal transduction, or *cis*/*trans*-regulatory mechanisms [1,2]. Indeed, many QTLs that influence some degree of QDR have been identified [8,13,14]. Linkage of such QTLs or the underlying loci to exact resistance features, like the lag-phase duration, will be one of the future challenges that would allow understanding and utilizing QDR in pathogen resistance breeding.

### Phenotyping technology and approaches to quantify QDR

The functional characterization of QDR highly depends on precisely measured phenotypes [2,15]. However, the experimental design required to assess QDR phenotypes over entire plant or pathogen populations quickly exceeds the limits of traditional, manual scoring methods and calls for more sophisticated phenotyping technology. The increasing availability of sensor technology (e.g., RGB, multi- or hyperspectral sensors) and analytical methods (e.g., deep-learning or artificial intelligence algorithms) recently have strengthened the attention to plant phenotyping [16]. Many studies have shown how imaging technology can be used to determine plant phenotypes like plant height, nutritional status, or water-use efficiency but also to assist breeder’s decisions [14,17,18]. Moreover, several reviews recently summarized the potential of modern sensor technology and related software in quantifying phenotypes of host-parasite interactions on multiple levels [19–24]. Even advanced applications, like in-field phenotyping or assessing complex features in non-standardized conditions, are possible due to deep-learning models like ‘PLPNet’ or ‘ResNet-9’ [25–27]. However, large phenotyping platforms also have limitations. High-end systems often collect a multitude of 3D scanning images or images in multiple spectral wavelengths. Analysis of these data is computationally intensive and often requires very specific knowledge. Thus, such technologies might overwhelm (non-data-science-) researchers with high amounts of complex datasets as significant skills are required to derive easy-to-interpret insights relevant to answering biological research questions [28]. A second challenge lies in adapting an established phenotyping system for various pathosystems, i.e., different crops or pathogens [22,29]. Lastly, most high-end phenotyping systems have very high investment and running costs and thus are less available. Combined with the aforementioned low flexibility, this further limits their use and application in the broad spectrum of plant pathology, where quick and easy screening of QDR in a large panel of plants is one of the main objectives. Recent developments, however, enable researchers to use the generally available consumer-level technology and build low-cost phenotyping platforms like the ‘Navautron’ [30]. In this study, we show the usefulness of such systems in unraveling QDR dynamics in crop wild relatives.

### Wild tomato populations as a reservoir of potential QDR loci against major pathogens

The domestic tomato (*Solanum lycopersicum*) is a major food crop of global importance [31]. However, plant pathogens, including the necrotroph *Sclerotinia sclerotiorum* or species from the genus *Alternaria,* commonly threaten tomato production worldwide [32–35]. Host resistance and fungicides are the standard tools to protect tomatoes against these pathogens. However, strong bottleneck events caused by R-genes or fungicides and higher-than-expected pathogen diversity in the field result in losing fungicide efficacy or plant resistance against such species [36–40]. Therefore, highly diverse wild populations are an invaluable source of desirable alleles in breeding, as crosses between wild and domestic lead to increased performance and stress tolerance [41]. Integrating phenotyping with screening of genetically highly diverse wild resources will help characterize novel alleles for QDR breeding [42].

Wild tomato species originated from several radiation events and can generally be classified into four groups within the so-called section Lycopersicon, containing a total of 15 species and two species in the section Lycopersicoides [43]. All species have adapted to specific habitats ranging from the edge of the Atacama desert to the Andes, where they withstand diverse (a)biotic stresses. Evolutionary analyses show that different species and populations have evolved drought or salt stress tolerance, as well as adaptation to cold stress [44–48]. Previous studies have also shown substantial variation in susceptibility and resistance of wild *Solanum* spp. against various pathogens but often relied on manual or single time-point disease assessments, thus lacking the temporal resolution and statistical power to describe QDR strategies confidently [38,49,50]. In light of the variation of QDR already shown, wild tomato species are perfectly suited for quantification of QDR mechanisms as proof of principle. Moreover, defining whether specific QDR mechanisms play major roles in resistance will generate much-needed insights into the biology of QDR to help design future durable resistance breeding projects against major pathogens.

*Sclerotinia sclerotiorum* is a necrotrophic pathogen that can infect hundreds of host species, including important crops such as rape seed and tomato [30,51,52]. On vegetables, including tomatoes, infection with *S. sclerotiorum* can cause tremendous yield loss due to collapsing stems or damaged fruits [53,54]. Infection in the field can happen through air-dispersed ascospores or via myceliogenic germination of its overwintering structures in the soil, the so-called sclerotia [32,52]. In experimental conditions, mycelial inoculation procedures are commonly used, as the preparation of ascospores can display a major challenge [55–59]. No complete form of resistance against the generalist *S. sclerotiorum* has been characterized; therefore, resistance breeding relies on QDR as the source of new alleles [32,52,55,57,60].

In the present work, we build on a low-budget image-based phenotyping system [30] to derive high-resolution time-resolved disease phenotypes and dissect them into three distinct QDR strategies. We show the potential of this system by characterizing the natural diversity of QDR phenotypes of wild *Solanum* species and, therefore, provide insights into the mechanisms underlying QDR against the generalist pathogen *Sclerotinia sclerotiorum*. We use this system as a model to address whether QDR is always represented by a similar mechanism, i.e., infection frequency or lag phase duration, and show that the orchestration of different QDR mechanisms affects the overall QDR on a genotype-specific basis. Accordingly, we argue that the different host species have evolved specific strategies to maintain a defined degree of QDR.

## 2. Materials and Methods

### Experimental Design

We screened multiple accessions of four wild tomato species (*S. pennellii, S. lycopersicoides, S. habrochaites,* and *S. lycopersicoides*) with a detached-leaf assay. All accessions of the same species were tested as one batch for up to five independent repetitions. To facilitate comparability between batches, *S. lycopersicum* cv. C32 was used as a control in every experiment. A schematic of the experimental procedures is displayed in fig. 1.

**Figure 1:**
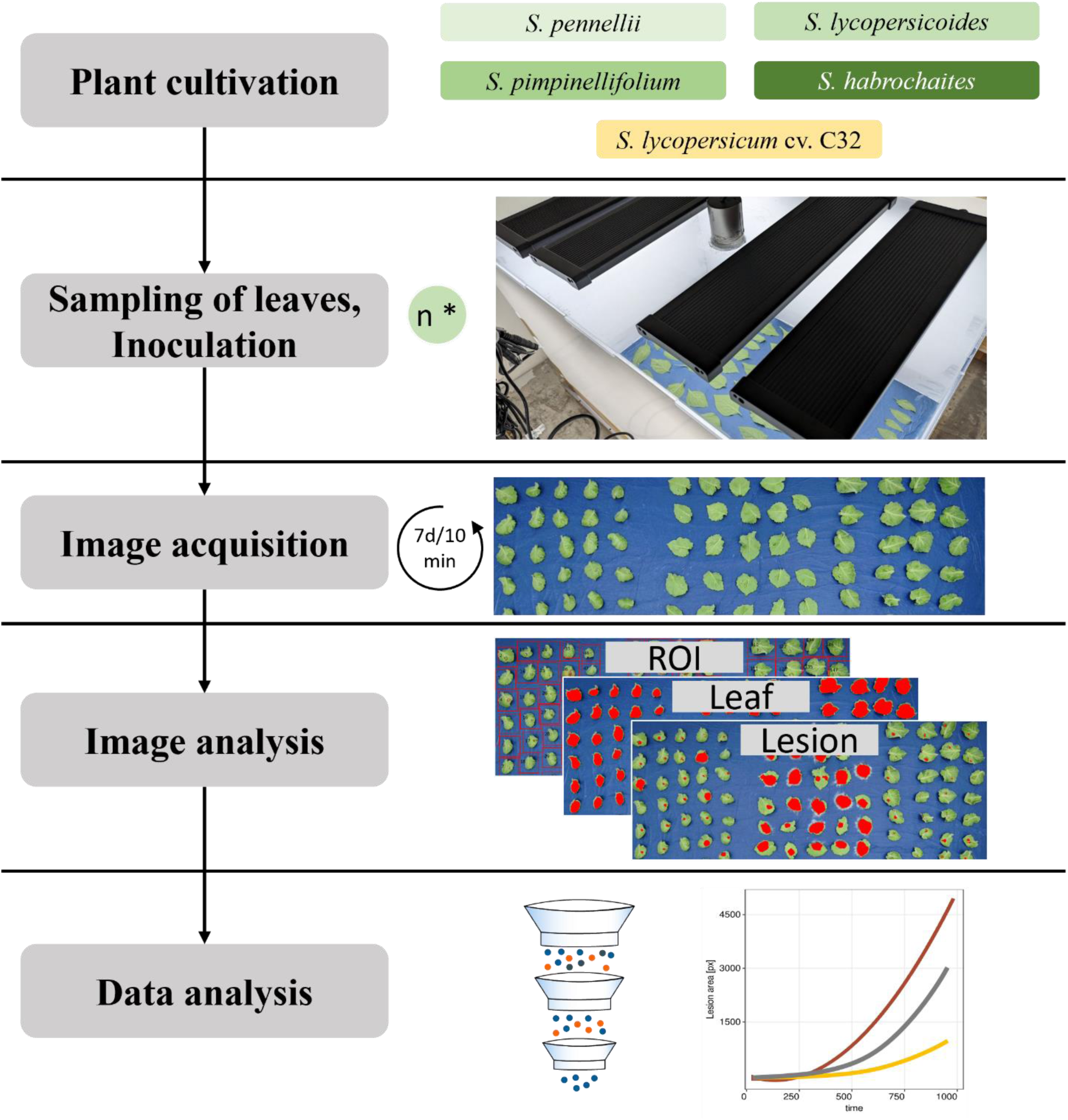
Overview of the high-throughput phenotyping assay.

### Sclerotinia sclerotiorum inoculum preparation

For inoculation experiments, the *Sclerotinia sclerotiorum* isolate 1980 or the OAH1:GFP isolate (for microscopical analysis only, [61]) was used. The fungus was alternatingly cultivated on potato dextrose agar (Sigma Aldrich) and solid malic acid medium [62] at approx. 25°C in the dark. Four 1cm pieces of *S. sclerotiorum* inoculum were used to inoculate 100 mL PDB. After four days of incubation on a rotary shaker (24°C, 120rpm), a fungal mycelium suspension was generated: for this, the medium was mixed using a dispenser (IKA T25) for two times 10 sec at 24.000 rpm. The mixture was then vacuum-filtrated through cheesecloth, and the remaining liquid was concentrated to an OD of 1. For the negative control, fungal tissue was removed from the solution by centrifugation, and the supernatant was autoclaved. Tween20 was used as a surfactant. Per leaf, one drop (10 µL) of inoculum was used.

### Plant growing conditions

Wild tomato germplasm was obtained from the C. M. Rick Tomato Genetics Resource Center of the University of California, Davis (TGRC UC-Davis, http://tgrc.ucdavis.edu/) (see suppl. table 5). The species were selected to include genetically diverse species within the section Lycopersicon and a species from the section Lycopersicoides (fig. 2). All plants were grown at the greenhouse facility of the Department of Phytopathology and Crop Protection, Institute of Phytopathology, Faculty of Agricultural and Nutritional Sciences, Christian Albrechts University, Kiel, Germany. Following seed surface sterilization using 2.75% hypochlorite (15 min. incubation followed by washing twice with dH_2_O), seeds were sown in the substrate (STENDER C700, Germany) and cultivated in a growth chamber (21 °C, 65% rH, 450 PAR). From the 3-leaf stage on, plants were cultivated in standard greenhouse conditions with supplement light. Plants were occasionally fertilized via the irrigation system (1% Sagaphos Blue, Germany). Plants were propagated using cuttings (Chryzotop Grün 0.25%) and regularly screened for virus infection.

**Figure 2:**
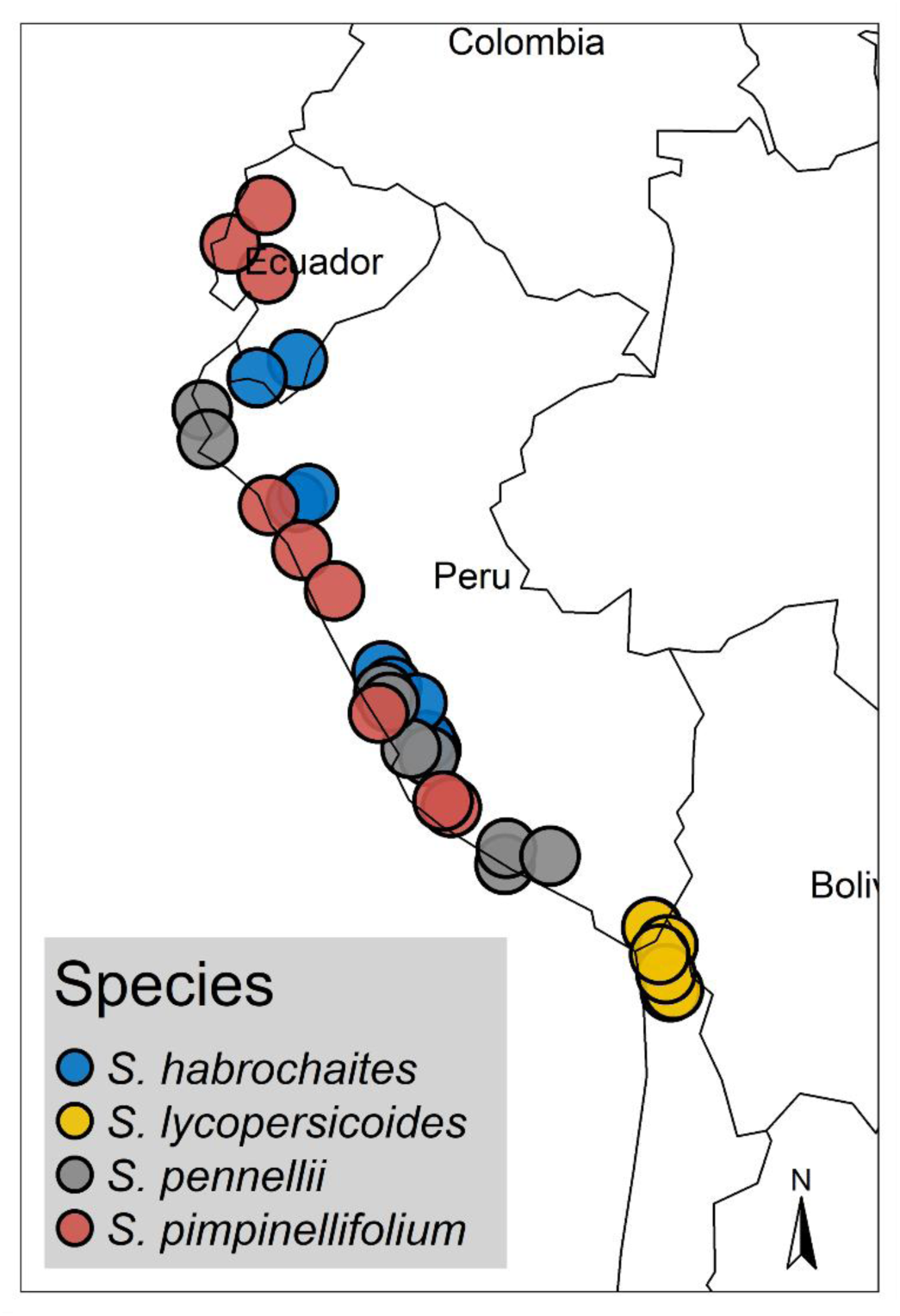
Sampling localities of wild tomato accessions used in this study. Seed material of all wild tomato accessions was provided from C. M. Rick Tomato Genetics Resource Center of the University of California, Davis (TGRC UC-Davis, http://tgrc.ucdavis.edu/). Individual dots represent the geographical origin of each accession.

### Detached leaf assay

Detached leaf assays were conducted to measure quantitative disease resistance of a diverse panel of wild Solanaceae plants. A custom phenotyping system was adapted [30]. A 50cm x 70cm PMMA tray was filled with eight layers of blue tissue paper and flooded with 700mL sterile dH2O. Plant leaves were placed abaxial side up onto the tissue and inoculated with 10 µL of mock-/*S. sclerotiorum-*suspension. Next, the tray is covered with a custom hood. The boxes were placed inside a growth chamber (24°C) and incubated for seven days. The assay was independently repeated five times. We used a representative set of three experiments for all further analysis.

### Phenotyping platform

High-resolution images were acquired using RGB cameras (Yealink UVC30) mounted on the box. Cameras were controlled using Raspberry Pi microcomputers or desktop PCs running headless Ubuntu22. A cron daemon launched the image-acquisition script every ten minutes. Plant lights also briefly illuminate during nighttime for image capture to enable images in the dark while maintaining circadian rhythm. This was achieved by using the ‘Shelly Plus Plug S’ wifi plug.

### Image analysis

We adapted the ‘navautron’ software package (https://github.com/A02l01/Navautron). The image analysis involved manually defining regions of interest (ROI) using ImageJ (ImageJ Version 1.530). Further, HSV thresholds were optimized individually per box. For this, ‘assess_noChl.py’ was used, and an overlay was generated in Gimp (Version 2.10). Once binary masks represented the respective feature classes (leaf_healthy, leaf_diseased, background), the whole dataset was evaluated using the ‘infest.py’ script. Segmentation was iterated and classified pixel counted. The analysis includes functions from the python3 (Version 3.11.4) libraries ‘numpy’ (Version 1.25.2), ‘opencv’ (Version 4.8.1.78), ‘plantcv’ (Version 4.0.1), and ‘scikit-image’ (Version 0.22.0). The plantcv function ‘dilate’ was used to remove leaf edges containing shadows with ksize=9, i=1 [63]. To improve thresholding accuracy (e.g., filling holes) on the lesion, an index filter was applied [ndimage.generic_filter(mask, threshold, size=3, mode=’constant’)] with a condition to overwrite pixels deviating from the value of the majority of the surrounding pixels. np.sum(mask) was used to quantify the number of pixels in each feature class (lesion and leaf). Code and scripts can be found at https://github.com/seveein/QDR_Wild_Tomatoes.

### Microscopy analysis

Plant leaves were harvested and inoculated under standard conditions as described before but with either a GFP-expressing *S. sclerotiorum* strain, the *S. sclerotiorum* wildtype 1980, or the mock suspension. The leaves were evaluated at 12-hour intervals using a Zeiss Discovery V20 stereomicroscope under bright light and fluorescent illumination (Zeiss HXP120). Images were taken using an AxioCam MRc camera.

### Statistical analysis

An interactive R-script (R-Version 4.3.2, R-Studio 2023.12.1+402) was facilitated to extract lag-phase duration and LDT to quantify resistance characteristics [30]. Each leaf’s lesion size over time was fitted against a 4-degree polynomial regression. The fit to the measured data point was reviewed for each sample. Lesion doubling time (LDT) and lag phase were determined based on a segmented regression analysis, expecting two linear phases. First, a linear phase during the lag period (no symptom development), and second, a linear log growth (symptom) during the exponential growth phase. The start of exp(LDT) is considered the lag phase, while the LDT represents the log(slope) of the linear growing curve in this area.

A two-tier filtering pipeline was developed to increase accuracy and remove artifacts from the dataset. First, single time points with an arbitrary high/low leaf area were filtered per leaf. The top and lowest 2.5 % of the data points were trimmed for this. Next, individual leaves with unexpectedly high variability in leaf area were excluded from the data set. Therefore, samples with sd(leaf) > 10% of the mean(leaf) were removed from the dataset using a simple tidyverse (v. 2.0.0) pipeline.

As a measure of symptom development over time, the area under the disease progress curve (AUDPC) was calculated using the R-package agricolae (v. 1.3-6). General statistical analysis and visualization were conducted in RStudio (R-Version 4.3.2, R-Studio 2023.12.1+402 [64]), and the packages tidyverse [65], ggplot2 [66], ggpubr [67] and agricolae [68]. AUDPC is defined with *i*=time and *y_i_*=symptom severity at time=*i* as: [68]

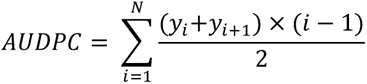

For continuous variables (lag phase duration, lesion doubling time, AUDPC, tt100), a statistical model based on a generalized least squares model was defined [69]. In contrast, a generalized linear model was defined for binomial values (infection frequency, 100%/f) [70]. These models included genotype and start date (without interaction effect).

The residuals corresponding to the continuous values were assumed to be approximately normally distributed and heteroscedastic concerning the different genotypes. These assumptions are based on a graphical residual analysis (suppl. fig. 7 & 8). Based on these models, a Pseudo R^2^ was calculated [71], and an analysis of variances (ANOVA) was conducted, followed by multiple contrast tests [72,73]. User-defined contrast matrices were used i) to compare the species’ means with each other and ii) to compare the population means within their specific species with the corresponding species’ mean. The individual leaf area was previously found to have no significant influence on lesion area; therefore, it was not included in our statistical model [74]. A linear mixed-effects model was used to determine the relationship between AUDPC and predictors such as genotype, lag phase duration, and LDT. Random intercepts were specified per start date to account for experimental repetitions.

Based on this model, fixed effect values were extracted and used to predict AUDPC per genotype_i=1,2,3_ in relationship to varying lag and LDT values.

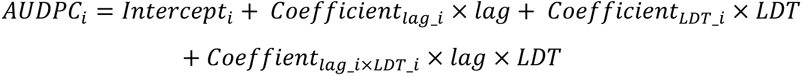

The associated R-codes can be found at https://github.com/seveein/QDR_Wild_Tomatoes.

## 3. Results

### Wild tomato species carry different levels of quantitative resistance against *Sclerotinia sclerotiorum* depending on defense-parameters

We investigated the phenotypic diversity in quantitative disease resistance in four wild tomato species (*S. habrochaites, S. lycopersicoides, S. pennellii,* and *S. pimpinellifolium*) against the *Sclerotinia sclerotiorum* isolate 1980 [13]. We used the “Navautron” automated phenotyping system for continuous image acquisition and applied a threshold-based segmentation algorithm to extract phenotypic data. Hence, we calculated different QDR parameters such as infection frequency, lag-phase duration, lesion doubling time (LDT), or area under the disease progress curve (AUDPC) to quantify temporal dynamics of infection (fig. 3). High variability between experimental runs with wild tomatoes has been described before [6,49,74]. To account for this, we applied a generalized least squares model (gls, continuous variables) and a generalized linear model (glm, discrete variables) for statistical analysis [69]. Overall, we discovered a great diversity of resistance phenotypes among the tested plant species. We found no 100% resistant accessions (suppl. fig. 3). We observed a significant difference in lag-phase duration among plant species, which we define as the time from infection until the first symptoms appear (see fig. 3 A, C, D). For instance, *S. pimpinellifolium* showed the shortest time from inoculation until lesion development (adjusted mean = 36.2 hrs). In contrast, *S. habrochaites* and *S. pennellii* displayed a significantly prolonged lag phase (both approx. 59 hours) (see suppl. table 1). Using segmented regression analysis, we determined the speed of lesion growth on individual leaves of the panel. The fastest-growing lesions were found on the species *S. pimpinellifolium* and *S. pennellii.* Lesions on *S. pennellii* and *S. pimpinellifolium* leaves doubled in size within approx. eleven hours (6.56 log(LDT) and 6.55 log(LDT), respectively), while lesions on *S. habrochaites* and *S. lycopersicoides* spread significantly slower. Those lesions expanded with an average rate of approximately 7.7 log(LDT), corresponding to roughly 36 (*S. habrochaites)* and 41 hours (*S. lycopersicoides)*(see suppl. table 2). Moreover, we observed that the success of disease establishment (infection frequency) depends highly on the host species. We identified a significantly lower infection rate on *S. habrochaites* (corrected infection frequency estimate 80 %), whereas *S. lycopersicoides* and *S. pennellii* displayed significantly higher infection frequency (∼93 and 95%, respectively) (fig. 3 B).

**Figure 3:**
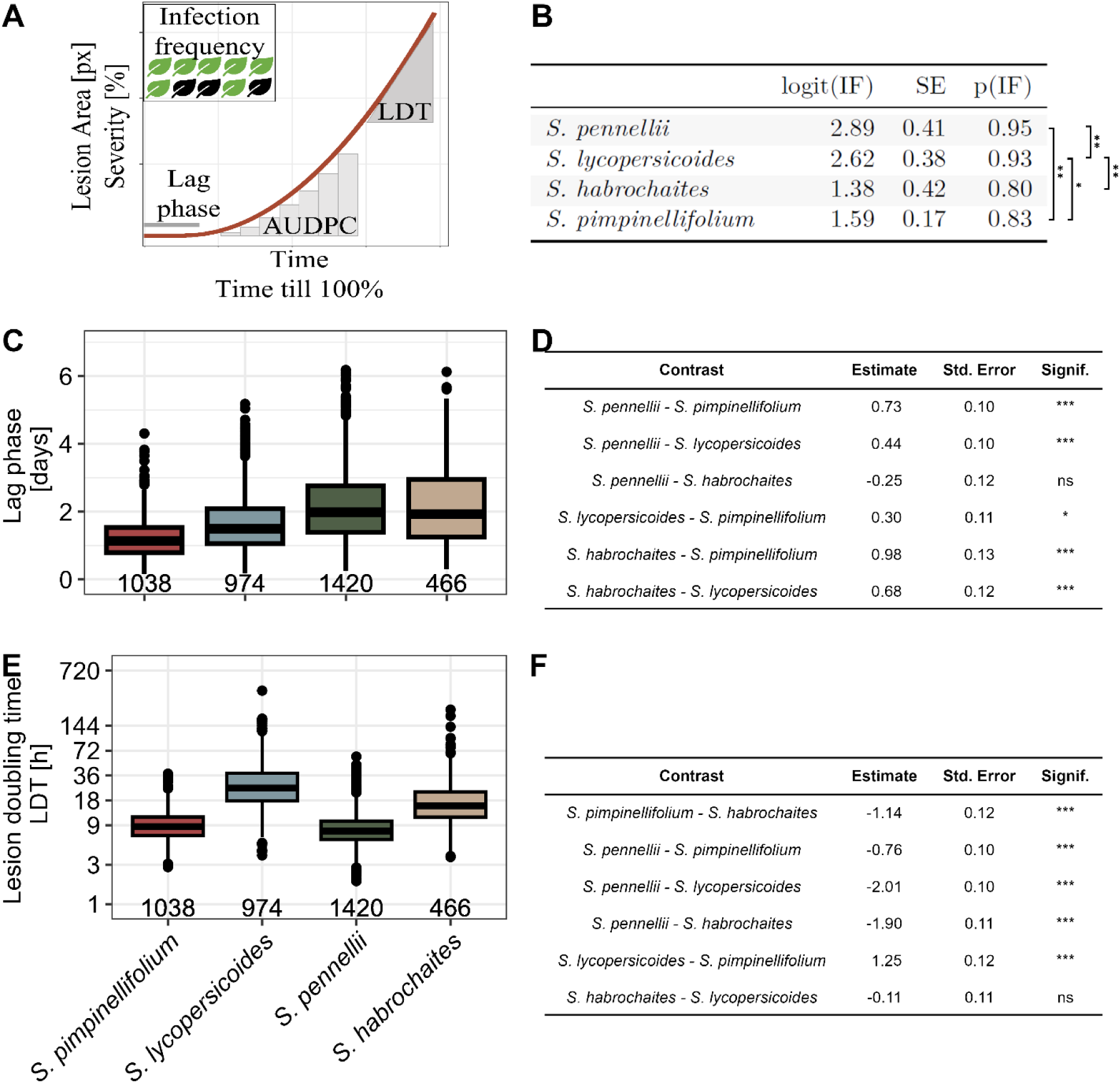
Wild tomato species possess a broad diversity of resistance against *S. sclerotiorum*. A) Exemplary illustration of different QDR parameters used in this study. The infection frequency is defined as the percentage of leaves showing a lesion after seven days of incubation. The lag phase, or lag-phase, is defined as the time till first visual symptoms appear. We used a segmented regression analysis to determine the lag phase’s end mathematically. The absolute lesion size is represented as pixel counts, whereas normalization against leaf area results in symptom severity. The area under the disease progress curve (AUDPC) is defined as the integral area under the severity curve, which depicts the severity over time. As a measure of the lesion spread, the Lesion Doubling Time (LDT) describes the time till a lesion doubles its size. The time till a lesion covers 100% of a leaf is described by tt100%. B) The infection frequency of *S. sclerotiorum* inoculum differs significantly between the host species. The table shows a meta-analysis of pooled accessions collected from three independent experiments. C) Time till lesion formation (in days). The number on the x-axis indicates the count of individual leaves tested. D) Statistical analysis of pairwise differences in lag phase duration between the tested wild tomato species. Values are displayed in days and derived from a generalized least squares model. E) Lesion growth rate during the exponential growth phase hours, plotted on a log scale. The number on the x-axis indicates the count of individual leaves tested. Raw values are plotted. F) Statistical analysis of pairwise differences between the tested wild tomato species regarding lesion doubling time. Values are displayed as log(LDT[h]). Levels of significance are displayed as *** P < 0.001, ** P <0.01. * P<0.05, P<0.1.

### Individual QDR measures show different levels of intraspecific variation and conservation on *S. pennellii* and *S. lycopersicoides* accessions

To assess the within-species diversity of QDR phenotypes, we tested different accessions of each represented species. We collected phenotypic data from seven *S. lycopersicoides* and nine *S. pennellii* populations (fig. 4), as well as eight populations of *S. habrochaites* and ten of the species *S. pimpinellifolium* (see suppl. fig 1 and suppl. fig. 2). Especially the comparison of *S. lycopersicoides* and *S. pennellii* highlights that QDR diversity differs between species. We observed that the (adjusted) mean duration of the lag phase on different *S. pennellii* accessions ranged from 1.59 days (38 hours, LA1809) to 2.86 days (68 hours, LA1303) (fig. 4 A, C). Using a generalized least squares model, we identified accessions with a significantly shorter lag phase than the grand mean of the species (LA1809 and LA2657). In contrast, the accessions LA1656 and LA1303 displayed a significantly longer lag phase (2.75 days [66 h] and 2.86 days [68 h], respectively) (fig. 4 A, C). Next, we observed a significantly shorter overall lag phase duration of *S. lycopersicoides* accessions than *S. pennellii.* Accordingly, the first symptoms appeared after 1.3 days (31 hours, LA2772) and the latest at 1.83 days after inoculation (43 hours LA1966). The overall time till initial symptom development was more conserved; only two *S. lycopersicoides* accessions deviated significantly from the grand mean, being more susceptible than the overall species level (LA2776 and LA2772) (fig. 4 B, D). Similarly, we found a lack of variation in lag-phase-duration in the populations of *S. pimpinellifolium*. At the same time, *S. habrochaites*-accessions displayed a wider variability of lag-phase phenotypes (suppl. fig. 1, suppl. table 3).

**Figure 4:**
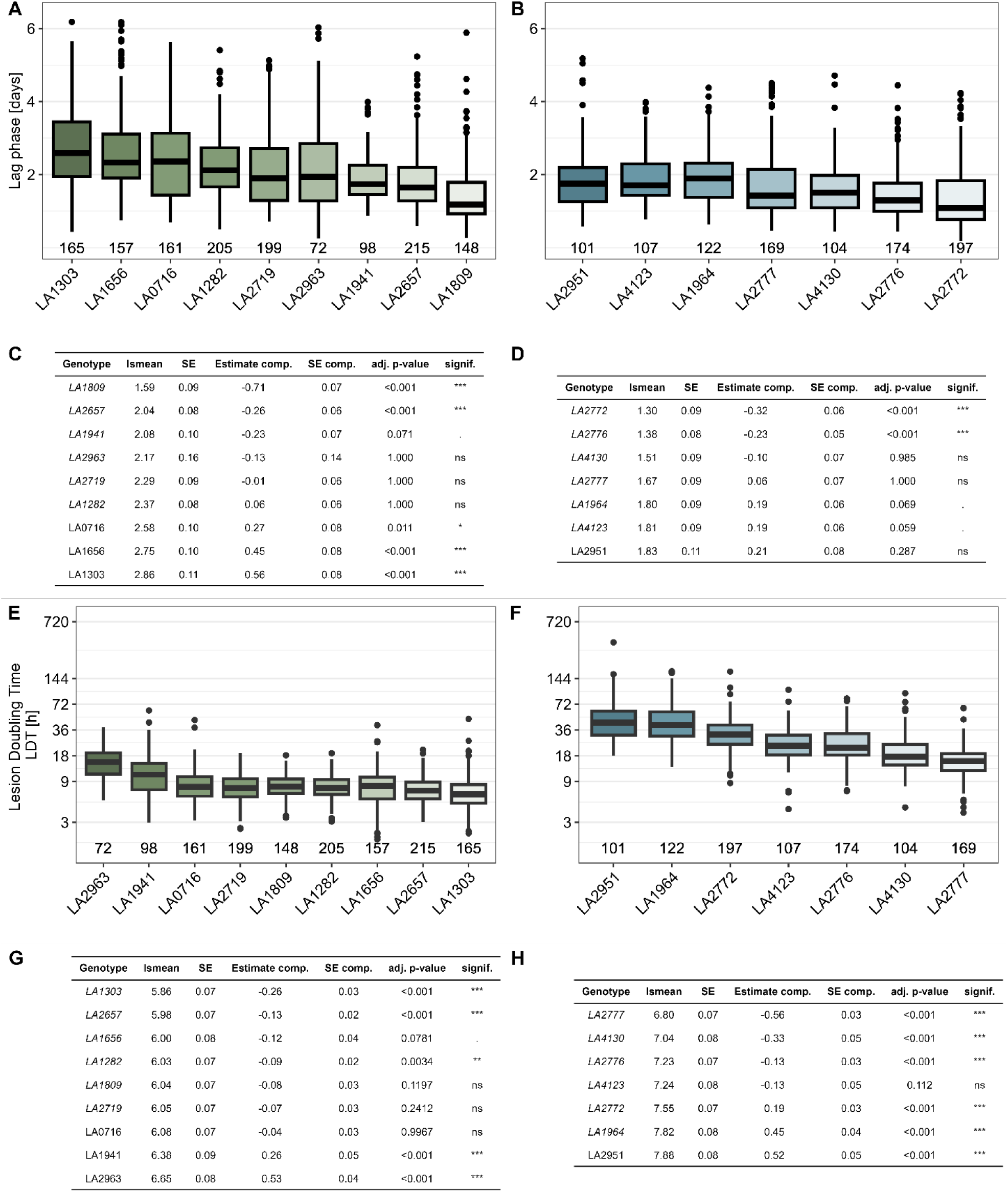
QDR parameters show different levels of variation depending on the host species (*S. pennellii and S. lycopersicoides)*. A) The lag phase duration (in days after infection) of *S. sclerotiorum* infection on *S. pennellii* accessions displays a higher level of intraspecific diversity than on accessions of *S. lycopersicoides* (B). C) & D) Variation statistics of the lag phase duration contrasting each accession with the grand mean per species (*S. pennellii* C, *S. lycopersicoides* D). Estimates are displayed in days post inoculation. E) & F) The lesion doubling time (in hours) of *S. sclerotiorum* infection on *S. pennellii* accessions is lower than on *S. lycopersicoides*. G) Variation statistics of LDT on *S. pennellii* and H) *S. lycopersicoides.* Lsmean-values and SE indicate the adjusted mean and SE per population. Estimate comp., SE.comp, and p-values describe pairwise statistics of each accession against the grand mean. The numbers on the x-axis in panels A, B, E, and F indicate the count of individual leaves tested. Levels of significance are displayed as *** P < 0.001, ** P <0.01. * P<0.05, P<0.1.

Next, we analyzed the variability of the lesion growth rate between accessions of each species using the logarithmic lesion doubling time. We observed that all tested *S. pennellii* accessions displayed an average lesion doubling time ranging from 5.84 h (LA1303) to 13.07 hours (LA2963). Five accessions (LA1809, LA1282, LA2719, LA2657, LA1303) have a significantly faster lesion development than the grand mean (LDT < 11 hours). The populations LA2963 and LA1941 displayed a significantly longer LDT (13.07 and 9.8 hours, respectively) (fig. 4 E, G). Generally, we found that symptoms of *S. lycopersicoides* grew significantly slower (observed range: 14.9 h to 40 hours). However, we still observed a significant within-species variability. For instance, symptoms on leaves of the accession LA2951 doubled within lsmean=7.88 log(LDT) (approx. 44 hours), while lesions of LA2777 expanded much faster at lsmean=6.8 log(LDT) (15 hours, fig. 4 F, H). We observed a high variability among the accessions for *S. pennellii* and *S. lycopersicoides*, mostly deviating from the species-mean in LDT with high significance. Interestingly, the LDT on *S. habrochaites* and *S. pimpinellifolium* appeared much more conserved between the accessions, as only a few samples significantly differed from the grand mean (suppl. fig. 2, suppl. table 4).

### Disease resistance measures are not linked and characterize distinct components of quantitative disease resistance

To test whether fungal infection is directly linked to lesion growth, we conducted microscopy assays using a GFP-tagged *S. sclerotiorum* mutant of the *S. sclerotiorum* isolate 1980 [75]. We selected two accessions from *S. pennellii* with significantly altered lag-phase duration. At 72 hours after inoculation, freshly developed mycelium was observed on leaves of the *S. pennellii* accession with the shortest lag phase duration (LA1809). In contrast, on the less susceptible accession LA1303, the first fungal structures started growing at 96 hpi (suppl. fig. 6). Fluorescent microscopy imaging showed that fungal mycelial structures were always accompanied by clear formation of necrotic lesions but cannot be observed prior visual lesion development (fig. 5 C, suppl. fig. 6). Thus, showing that a longer lag phase does not represent any latent or biotrophic infection and that LDT and lag phase are likely uncoupled phenomena.

**Figure 5:**
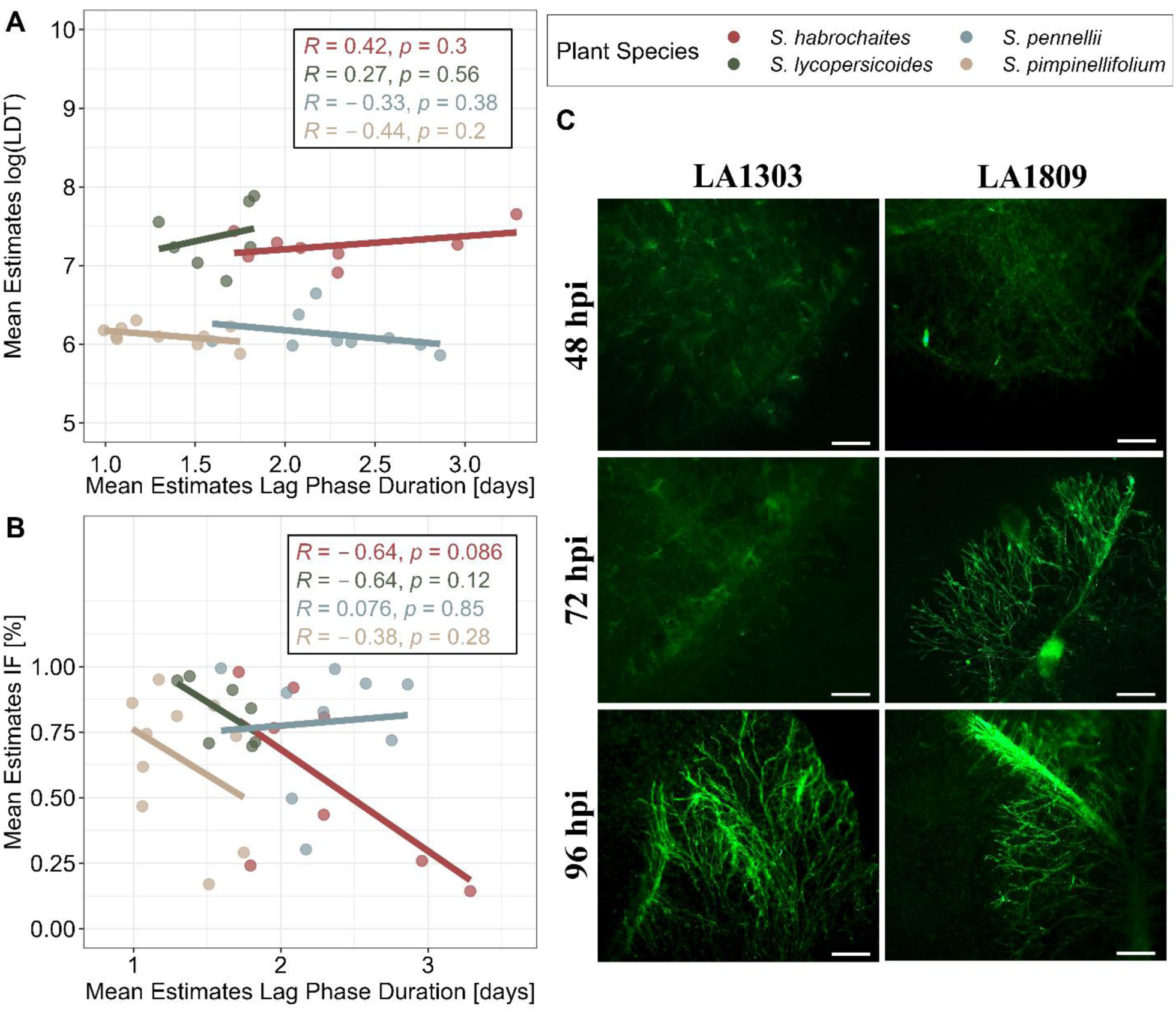
Different QDR parameters are independent from each other. A) Pearson correlation analysis of LDT and lag-phase duration. Dots represent the least squares of each accession. B) Pearson correlation analysis between the infection frequency and lag phase duration. Dots represent infection frequency adjusted per-accession estimates from glm/gls. C) The lag-phase duration of *Ss1980:GFP* infection on *S. pennellii* genotypes is reflected in fungal growth dynamics. Images show a representative selection of 10 biological replicates. The bar indicates 500 µm.

We performed a correlation analysis to consolidate the relationship between the QDR parameters further. First, we tested the overall relation of lsmean LDT and lsmean lag-phase duration by pooling all accessions of all species. We found that LDT and lag phase were independent (R=0.14), with no significant relationship (p =0.42) (suppl. fig. 4). We also tested the correlation between QDR strategies at the species level. We found only minor linear relationships between LDT and lag-phase for the four tested species. However, we found a weak, significant negative correlation between infection frequency and the duration of the lag phase (lsmean) in *S. habrochaites* (R = - 0.64, p=0.086) (fig. 5 A, B). For the remaining species, no significant correlation was found. We did not find a single host accession with high levels of resistance in both, LDT and lag-phase duration.

### Severity analysis reveals distinct resistance phenotypes against *Sclerotinia sclerotiorum* within a single species

For an in-depth analysis of disease severity, we selected three *S. pennellii* accessions with similar leaf sizes: LA1282, LA1809, and LA1941 (fig. 6 A). While symptoms developed on most of the leaves, the impact of infection is highly dependent on the respective accession (see fig. 6 B). Accession LA1941 shows a significantly lower infection frequency (∼51%) and a significantly lower rate of fully infected leaves than LA1809 (approx. 11% vs. approx. 41%) or LA1282 (approx. 33%, fig. 6 C). It took approx. 6.5 -7 days to cover the whole leaf surface of LA1282 and LA1941. We found a significantly faster lesion spread on LA1809 with approx. 5.5 days till 100 % coverage.

**Figure 6:**
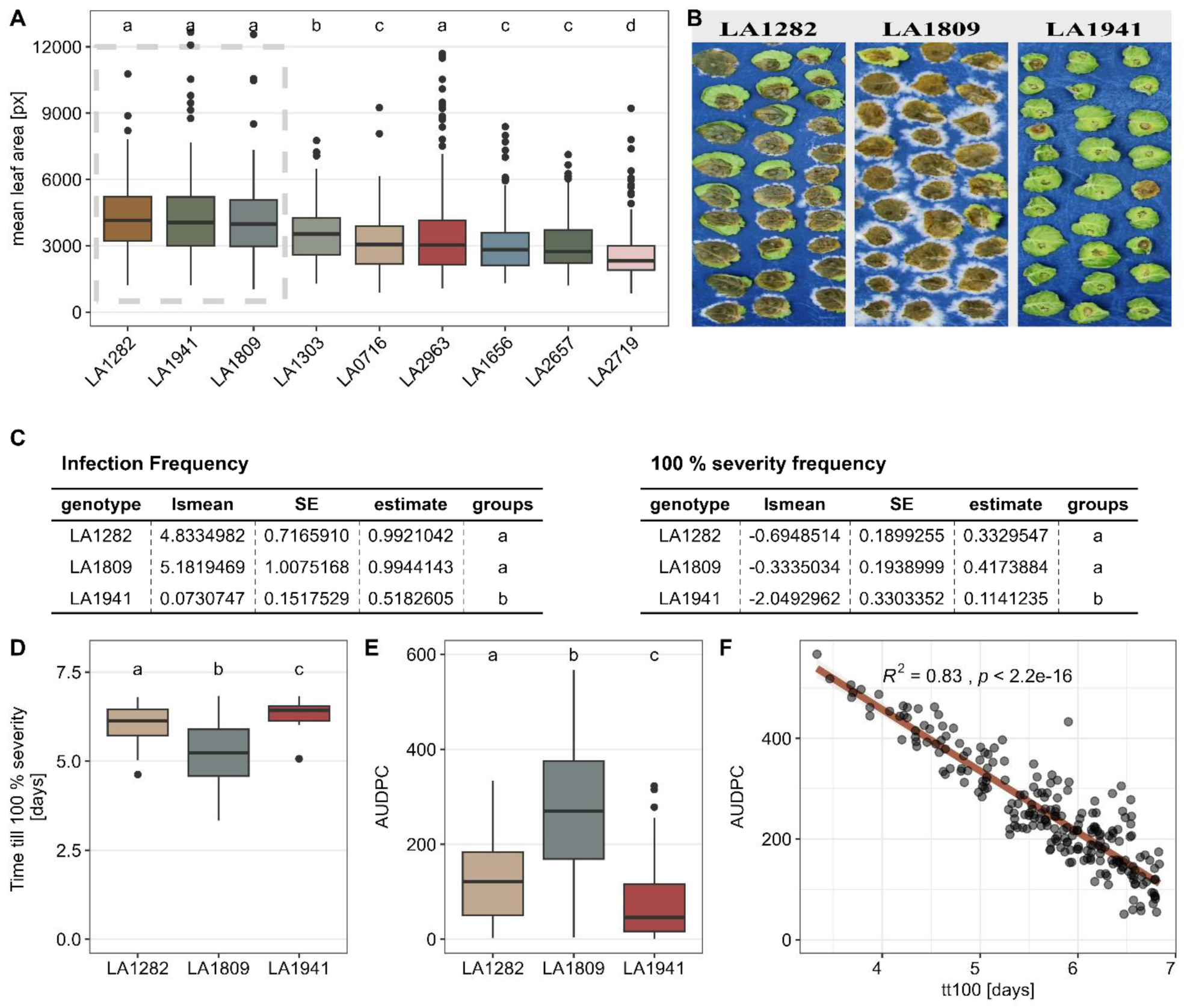
*S. pennellii* accession LA1941 harbours significantly elevated level of quantitative resistance against *S. sclerotiorum*. A) Mean leaf area of *.S. pennellii* accessions quantified during infection experiments. The data of three independent experiments is shown. HSD test was performed to identify cluster with similar leaf size. Selected plants with similar leaf size are indicated by the box. B) Exemplary images of *S. sclerotiorum* infections on the *S. pennellii* populations LA1809 and LA1941 at seven days post-inoculation. C) Statistical analysis of Infection frequency (IF) and frequency of fully infected leaves at the end of experiment. “lsmean” represents the estimate as logits, while ‘estimate’ represents the estimated probability. D) Comparison of time till lesion saturation of *S. sclerotiorum* on *S. pennellii* genotypes. E) Area under disease progress curve (AUDPC) of three *S. pennellii* populations with similar leaf size. Wilcoxon-test was performed for levels of significance. Time series data from previous experiments was used. F) Pearson correlation analysis of tt100 vs. AUDPC. All statistics were calculated using a glm/gls with custom contrast matrizes. Compact letter display were determined using the package ‘multcompLetters’ with a threshold of p < 0.05.

**Figure 7:**
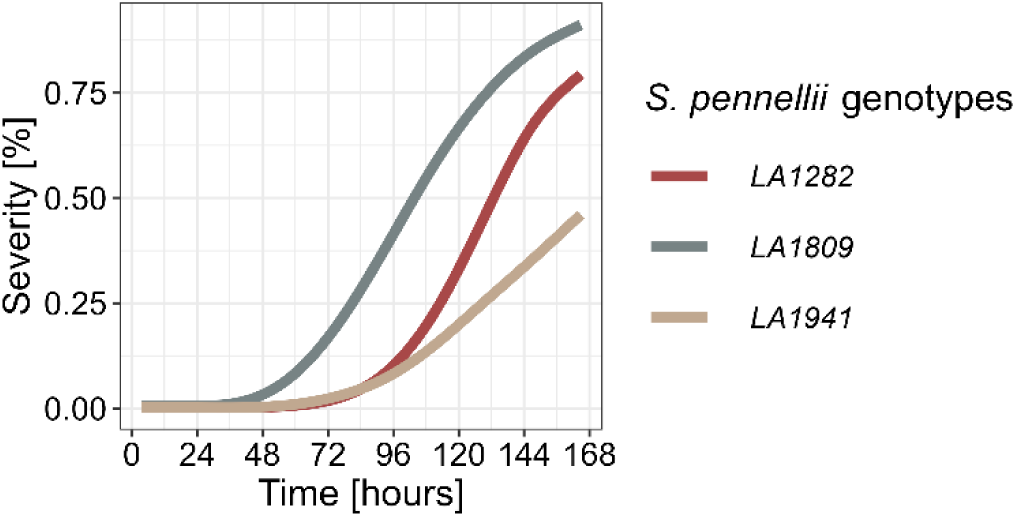
Exemplary growth curve of three *S. pennellii* accessions with different resistance levels against *S. sclerotiorum*. Shown is the mean symptom-severity of each accession as share of leaf area over the period of seven days. The experiment was independently repeated three times. n_LA1282_ =205, n_LA1809_=148, n_LA941_=98.

This is also reflected by the AUDPC, where LA1809 had the highest mean value (AUDPC approx. 250). In contrast, in LA1282 and LA1941, the symptoms spread much slower, leading to significantly lower AUDPC values (100 and 50, respectively, fig. 6 E). Together, the prolonged time till 100% severity and a considerably low AUDPC on LA1282 indicate a late but explosive lesion growth, corresponding to the mean severity-kinetics of those genotypes (fig. 6).

### The moderation of QDR parameters is genotype-dependent

Next, we used a linear mixed-effect model (lme) to test which of the factors have the strongest effects on disease severity on those accessions (*S. pennellii* LA1282, LA1809, LA1941). Following the analysis of variance (ANOVA), we found a significant influence of most tested variables (genotype, lag phase, LDT) on the AUDPC (tab. 1). Strikingly, we found that the genomic background of the tested plants is insufficient to explain the observed diversity in AUDPC. In other words, we observe a significant relationship between lag, LDT, and their interaction with the genotype. Because of this, we extracted the fixed-effect estimates from the lme and generated predictor functions for the AUDPC of each genotype. Then, we modeled the AUDPC using high-confidence lag and LDT values from previous observations (see fig. 3 C, D). We observed the highly variable influence of lag-phase duration, LDT, and their interaction on the AUDPC (fig. 8). Strikingly, we found that variation of the LDT has almost no influence on the AUDPC of LA1809 besides the generally elevated severity level (fig. 8). Further, we found that only a prolonged lag phase duration might contribute to an increased potential for lower severity in LA1809 (fig. 8). However, the influence of longer lag-phase is reduced with increasing LDT. For leaves of the accessions LA1282 and LA1941, we found a stronger combined effect of lag-phase and LDT on the severity. More specifically, a prolonged lag phase might lead to a small reduction of the symptom severity on LA1282 while reducing the AUDPC on LA1941 more rapidly. Further, we observe that a prolonged lesion doubling time reduced symptom severity in both LA1282 and LA1941.

**Figure 8:**
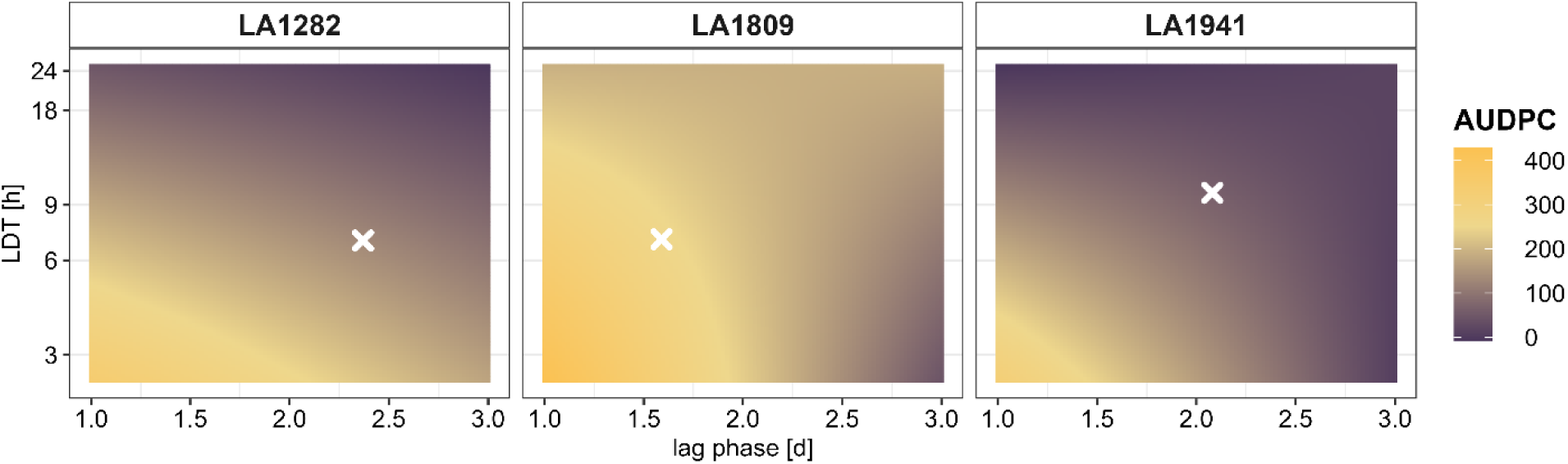
QDR parameters contribute highly dynamic and host-specific to symptom severity. We used the *S. pennellii* accessions LA1282, LA1809 and LA1941 to test for the genotype-dependent relationship between lag and LDT. Therefore, we extracted the estimates for the factors LDT and lag per each genotype from an ANOVA based on a generalized least square model (tab. 1). The per-genotype AUDPC was modeled using the extracted estimates over a range of values representing the plausible range of lag-/LDT-values. Crosses represent the observed mean AUDPC (fig. 6 E).

**Table 1:** Statistical analysis of the effects of genotype, lag-phase duration, LDT, and their interactions on disease severity (AUDPC) of the *S. pennellii* accessions LA1282, LA1809, and LA1941. Results of an analysis of variance (ANOVA) based on a linear mixed-effects model are shown.

|  | numDF | denDF | F.value | p.value |
| --- | --- | --- | --- | --- |
| Intercept | 1 | 437 | 136.8 | <0.001 |
| Genotype | 2 | 437 | 211.34 | <0.001 |
| Lag | 1 | 437 | 251.68 | <0.001 |
| LDT | 1 | 437 | 90.41 | <0.001 |
| Genotype:Lag | 2 | 437 | 8.54 | <0.001 |
| Genotype:LDT | 2 | 437 | 2.32 | 0.099 |
| Lag:LDT | 1 | 437 | 21.91 | <0.001 |
| Genotype:Lag:LDT | 2 | 437 | 3.3 | 0.038 |

## 4. Discussion

### QDR against *Sclerotinia sclerotiorum* is highly diverse in *Solanum* spp

Wild tomato species have been screened for quantitative resistance phenotypes against many diseases, including Tomato brown rugose fruit virus*, Phytophthora infestans, Alternaria solani, Fusarium spp.* or *Botrytis cinerea* [8,49,50,74,76–79]. However, high hurdles in characterizing QDR on a phenotypic level limit detailed insights into the functional role of QDR against necrotrophic pathogens. This was mostly due to the lack of affordable high-throughput phenotyping facilities [1,2,5,8,80]. Here, we present a unique dataset of high-resolution QDR phenotypes against *Sclerotinia sclerotiorum* on a diverse set of wild *Solanum* species derived from a low-budget phenotyping set-up. In total, we tested almost 7,000 leaves with approx. 1,000 measurements each, resulting in approx. 7 million data points. We used this unique dataset to characterize the lesion development of infected leaves and applied advanced statistical analysis methods to extract more specific descriptors for QDR, such as lag phase, LDT, or AUDPC [30]. Because of this system’s scale and temporal resolution, we generated novel insights into the phenomena contributing to QDR.

### Interspecific QDR phenotypes follow a wide distribution

As expected, we observed a diverse range of disease phenotypes, as demonstrated in previous studies[6,49,78]. None of the tested accessions carried complete resistance against *S. sclerotiorum,* although we found a wide distribution of infection phenotypes. Also, no high ‘universal’ level of partial resistance or tolerance among multiple QDR parameters was found, as none of the species harbors significant advantages in multiple measures (infection frequency, lag phase, or lesion doubling time). Complete resistance against *S. sclerotiorum* is rarely found in cultivated crops [32,52,60,81]. We provide evidence that the time till the emergence of the first lesions (lag-phase) is highly variable within and between host species, with only *S. lycopersicoides* showing a rather conserved lag-phase duration (fig. 4 B). Interestingly, Barbacci et al., (2020) reported that in *Arabidopsis thaliana* the lag-phase duration is mostly influenced by the *S. sclerotiorum* isolate rather than the host accession. The comparably low genetic diversity of the host may have influenced the observed range of QDRs. Standing genetic variation is considered much higher in (predominantly) outcrossing *Solanum* species than in inbreeding *A. thaliana* accessions [82]. Accordingly, we assume that the influence of genetic features on the lag-phase duration depends on the specific genomic background of the host plant species. However, fungal influences on pathogenesis cannot be ignored, as the concept of the ‘extended phenotype’, describing the interaction of both genomes i.e. a genotype x genotype (GxG) interaction for host and pathogen, for one phenotype, is well established [37,83]. Furthermore, quantitative host resistance features have been described to interact with the pathogen’s genotype as described for camalexin-associated resistance [50,84].

### QDR phenotypes also differ on the intraspecific level but at varying degrees

High variability of QDR phenotypes among genotypes of the same plant species has been reported on multiple hosts before [6,30,49,74,85,86]. We show that the degree of variability depends on the host species and the respective resistance parameter. Whereas LDT is rather stable among *S. pimpinellifolium* accessions, it is highly variable on *S. lycopersicoides* accessions (see fig. 4 & suppl. fig. 2). The specific forms of QDR phenotypes might hint at independent regulatory mechanisms and different evolutionary backgrounds with relatively recent developments, leading to genetic variation, rather than conserved QDR mechanisms. Host adaptation to natural habitats and its influence on disease resistance has been studied before [87,88]. Adaptation might explain disease phenotypes as most *S. lycopersicoides* accessions show significantly prolonged LDT. The habitat of *S. lycopersicoides* faces much more rain than the other species, leading to higher chances of successful infection events than in relatively dry habitats, thus requiring mechanisms to fight established infections. In contrast, drought-resistant *S. pennellii* has high capabilities in delaying infection events, while it lacks defense efficacy once an infection is established (fig. 3), similar to *S. chilense* desert population losing resistance against the fungus *Passalora fulva* [46,87,89]. However, to truly test these hypotheses, significantly higher sample sizes and infections under natural conditions would be required, possibly paired with screenings of the morphological properties of the species to assess the pleiotropic influence of habitat adaptation on QDR, e.g., via cuticle thickness or stomata density.

### QDR and Genotype x Genotype x Environment interactions

*S. pennellii* accession LA0716 was characterized as relatively resistant against *B. cinerea,* while this genotype is highly susceptible to *S. sclerotiorum* (suppl. fig. 5) [6]. Also in *S. chilense*, QDR phenotypes vary between the pathogen, suggesting the presence of pathogen-specific regulatory mechanisms [78]. However, the pathogen diversity tested in such studies might greatly affect the observed degree of resistance. A study with *Phytophthora infestans* on 85 *S. chilense* accessions showed that the relative differences in resistance phenotypes between individuals were mainly determined by the plant genotype, with modest effects of pathogen isolate used [49]. In contrast, large-scale screenings of infections with different *B. cinerea* isolates showed a clear genotype x genotype (G x G) effect both on panels of wild and domesticated tomatoes, as well as on *Arabidopsis thaliana* [37,50,74,90,91]. In addition, we have shown in *S chilense* that QDR phenotypes, like the infection frequency, can be correlated with the phytohormone ethylene [92]. Knowing that such phytohormonal regulation is also affected by abiotic, environmental (E) factors like temperature, humidity, and light availability, we propose that QDR polymorphism is implemented in a complex signaling network affected by GxGxE interactions [1,5,93,94].

### QDR is determined by the interplay of QDR strategies

QDR is commonly defined as a highly interconnected regulatory network with an integrated, pleiotropic role in general plant metabolism [1]. Therefore, the linkage of different defense strategies, like IF and lag-phase duration, could be a good perspective for resistance breeding. However, we did not observe strong correlations between QDR parameters and did not find a species or accession with a universal high resistance level for all tested parameters. Disconnected QDR parameters have been reported before: *Xanthomonas axanopodis* mutants showed increased infection frequency but a reduced lesion growth rate on cassava and *B. cinerea* showed unconnected IF and lesion expansion rates on wild tomatoes [6,95]. We used the presented phenotyping platform to show that the moderation or cross-talk between defense strategies is genotype-specific and differs even between accessions of the same species (fig. 8). Based on these findings, we propose a model for QDR against necrotrophic pathogens involving three genetically distinct strategies: 1. Prohibition of initial infection, 2. Retardation of disease outbreaks, and 3. Deceleration of ongoing infections.

### Disease severity is specifically determined by genotype-dependent moderation of QDR strategies

We used three differently severely infected *S. pennellii* genotypes to describe the influence of two of the QDR strategies (retardation and deceleration of symptom development) on overall symptom severity. Interestingly, the different accessions possess diverse capabilities in moderating the QDR strategies, as our model-based approach indicates contrasting roles of LDT and lag-phase duration.

In *A. thaliana,* it was shown that lesion traits, like lesion size or shape, are also controlled by genetically distinct mechanisms [90]. Previous work showed that defense-associated hormone responses greatly differ between different wild tomato accessions and even within the same population. In *S. chilense,* ethylene responses could only be linked to IF in one population but not in others [92]. Therefore, we argue that the genetic finetuning of QDR measures highly depends on the specific genetic background, and future studies should determine the complex interplay between various QDR-regulating strategies [94].

In this study, we used a new phenotyping platform to derive different QDR-related phenotypes. The low cost and high flexibility of the system allowed us to screen a big set of diverse plants relatively fast, and therefore, we identified new genotypes with distinct QDR properties. Accordingly, we characterized accessions and species with beneficial properties as significantly longer lag-phase duration (*S. pennellii,* LA1303 & LA1656) or prolonged LDT (*S. lycopersicoides,* LA2951 & LA1964). Accordingly, we suggest that *S. pennellii* accessions are specialized in delaying lesion development, whereas *S. lycopersicoides* accessions are more capable of slowing down the spread of established lesions. Follow-up research is needed to identify the genes underlying these differences. The resolution of the present dataset will enhance the ability to predict distinct defense phases, facilitating more targeted sampling strategies for transcriptomic or metabolomic analysis. This can help breed durable resistance in tomato crops with delayed and less severe symptoms without inducing strong evolutionary pressure. The sustainability of major R gene-mediated resistance (including pyramiding of such) has regularly been questioned [4,96]. Facilitating the concept of QDR is proposed to thwart the arms race between plant hosts and pathogens. QDR phenotypes specifically tolerate disease to a certain extent without applying a strong bottleneck onto the pathogen population [14]. Our findings provide major insights into the architecture of QDR strategies and will help in the targeted functional characterization of QDR. By disentangling end-point QDR phenotypes into discrete resistance mechanisms, the functional characterization of genetic features controlling QDR will become much more targeted. Based on this study, the factors influencing the level of QDR can be explained in much more detail.

## Acknowledgments

We gratefully acknowledge Hendrik Seide, Lilly Gieseler, and Ellen Krohn for their technical support and Dr. Farooq Ahmad for his insightful feedback and discussions whilst writing the manuscript. We also thank the TGRC at UC Davis (USA) for seed material.

## Availability of data and materials

Additional data can be found in the supplementary information files. All scripts used for this study are available at https://github.com/seveein/QDR_Wild_Tomatoes.

## Competing Interest declaration

The authors declare that they have no competing interests.

## Funding

This work was partly funded by the DFG (STA1547/6 and SFB924) and the ANR (ANR-21-CE20-30) for the collaborative project ResiDEvo. Exchange visits were supported by the DAAD and the Partenariat Hubert Curien programme of Campus France.

## Authors’ contribution

RS and SR conceived this study, SE planned and performed the experiments. AB and CTR developed the image analysis pipeline. SE conducted all analytical steps and produced the figures. MH developed the statistical tests and models. SE and RS wrote the manuscript. All authors read and approved the final manuscript.

## Notes

https://github.com/seveein/QDR_Wild_Tomatoes/

### Competing Interest Statement

The authors have declared no competing interest.

