## Supplementary Material for "High-resolution disease phenotyping reveals distinct resistance strategies of wild tomato crop wild relatives against *Sclerotinia sclerotiorum*"

##### Tables

Suppl. Table1: Estimate Means lag-phase duration

| Species | Lag estimate [h] | std.error [h] |
|---|---|---|
| ***S. habrochaites*** | 59.74 | 3.10 |
| ***S. lycopersicoides*** | 43.32 | 2.71 |
| ***S. pennellii*** | 59.87 | 2.71 |
| ***S. pimpinellifolium*** | 36.22 | 0.89 |

Suppl. Table 2: Estimate Means LDT

| Species | LDT-estimate [h] | std.error [h] |
|---|---|---|
| ***S. habrochaites*** | 36.85 | 0.02 |
| ***S. lycopersicoides*** | 41.13 | 0.02 |
| ***S. pennellii*** | 11.80 | 0.02 |
| ***S. pimpinellifolium*** | 11.77 | 0.02 |

Suppl. Table 3: Grand Mean contrasts lag

| Species | Accession | estimate (h) | std.error | statistic | adj.p.value | signif |
|---|---|---|---|---|---|---|
| ***S. habrochaites*** | LA1721 | -13.9805 | 3.2319 | -0.0721 | 0.0006 | *** |
| ***S. habrochaites*** | LA2167 | -12.0752 | 3.2768 | -0.0614 | 0.0078 | ** |
| ***S. habrochaites*** | LA1559 | -8.2475 | 2.3949 | -0.0574 | 0.0191 | * |
| ***S. habrochaites*** | LA1731 | -5.0888 | 3.3181 | -0.0256 | 0.9840 | ns |
| ***S. habrochaites*** | LA2128 | -0.1378 | 5.7960 | -0.0004 | 1.0000 | ns |
| ***S. habrochaites*** | LA2409 | -0.0536 | 2.5831 | -0.0003 | 1.0000 | ns |
| ***S. habrochaites*** | LA1753 | 15.8582 | 5.5225 | 0.0479 | 0.1275 | ns |
| ***S. habrochaites*** | LA2864 | 23.7252 | 6.5906 | 0.0600 | 0.0107 | * |
| ***S. pennellii*** | LA1809 | -17.0236 | 1.6687 | -0.1700 | 0.0000 | *** |
| ***S. pennellii*** | LA2657 | -6.3068 | 1.3466 | -0.0781 | 0.0001 | *** |
| ***S. pennellii*** | LA1941 | -5.4652 | 1.7853 | -0.0510 | 0.0713 | . |
| ***S. pennellii*** | LA2963 | -3.1692 | 3.2787 | -0.0161 | 1.0000 | ns |
| ***S. pennellii*** | LA2719 | -0.3337 | 1.5238 | -0.0036 | 1.0000 | ns |
| ***S. pennellii*** | LA1282 | 1.5361 | 1.3216 | 0.0194 | 0.9998 | ns |
| ***S. pennellii*** | LA0716 | 6.5780 | 1.8289 | 0.0599 | 0.0108 | * |
| ***S. pennellii*** | LA1656 | 10.7843 | 1.8850 | 0.0954 | 0.0000 | *** |
| ***S. pennellii*** | LA1303 | 13.4000 | 1.9487 | 0.1146 | 0.0000 | *** |
| ***S. lycopersicoides*** | LA2772 | -7.6031 | 1.5262 | -0.0830 | 0.0000 | *** |
| ***S. lycopersicoides*** | LA2776 | -5.5762 | 1.2054 | -0.0771 | 0.0001 | *** |
| ***S. lycopersicoides*** | LA4130 | -2.4261 | 1.5880 | -0.0255 | 0.9847 | ns |
| ***S. lycopersicoides*** | LA2777 | 1.4128 | 1.5828 | 0.0149 | 1.0000 | ns |
| ***S. lycopersicoides*** | LA1964 | 4.4474 | 1.4488 | 0.0512 | 0.0693 | . |
| ***S. lycopersicoides*** | LA4123 | 4.6261 | 1.4813 | 0.0520 | 0.0584 | . |
| ***S. lycopersicoides*** | LA2951 | 5.1190 | 1.9937 | 0.0428 | 0.2865 | ns |
| ***S. pimpinellifolium*** | LA1593 | -7.8207 | 0.8582 | -0.1519 | 0.0000 | *** |
| ***S. pimpinellifolium*** | LA2853 | -6.1720 | 1.5283 | -0.0673 | 0.0018 | ** |
| ***S. pimpinellifolium*** | LA1374 | -6.0821 | 1.1340 | -0.0894 | 0.0000 | *** |
| ***S. pimpinellifolium*** | LA1332 | -5.4925 | 1.1039 | -0.0829 | 0.0000 | *** |
| ***S. pimpinellifolium*** | LA1261 | -3.5010 | 1.6940 | -0.0344 | 0.7161 | ns |
| ***S. pimpinellifolium*** | LA1659 | -0.5469 | 0.8171 | -0.0112 | 1.0000 | ns |
| ***S. pimpinellifolium*** | LA4713 | 4.6669 | 4.1684 | 0.0187 | 0.9999 | ns |
| ***S. pimpinellifolium*** | LA2347 | 5.5052 | 1.0240 | 0.0896 | 0.0000 | *** |
| ***S. pimpinellifolium*** | LA1348 | 9.0943 | 1.1659 | 0.1300 | 0.0000 | *** |
| ***S. pimpinellifolium*** | LA2983 | 10.3488 | 1.5771 | 0.1094 | 0.0000 | *** |

Suppl. Table 4: Grand Mean contrasts LDT

| Species | Accession | Estimate | Std.Error | Statistic | Adj.p.value | signif. |
|---|---|---|---|---|---|---|
| ***S. habrochaites*** | LA2128 | -0.3450 | 0.0984 | -3.5052 | 0.0154 | * |
| ***S. habrochaites*** | LA2167 | -0.1418 | 0.0797 | -1.7798 | 0.9157 | ns |
| ***S. habrochaites*** | LA2409 | -0.1036 | 0.0609 | -1.7024 | 0.9466 | ns |
| ***S. habrochaites*** | LA1731 | -0.0322 | 0.0661 | -0.4866 | 1.0000 | ns |
| ***S. habrochaites*** | LA1753 | 0.0102 | 0.0890 | 0.1143 | 1.0000 | ns |
| ***S. habrochaites*** | LA1559 | 0.0349 | 0.0619 | 0.5633 | 1.0000 | ns |
| ***S. habrochaites*** | LA1721 | 0.1810 | 0.0837 | 2.1633 | 0.6332 | ns |
| ***S. habrochaites*** | LA2864 | 0.3965 | 0.1654 | 2.3970 | 0.4207 | ns |
| ***S. lycopersicoides*** | LA2777 | -0.5631 | 0.0319 | -17.6649 | 0.0000 | *** |
| ***S. lycopersicoides*** | LA4130 | -0.3304 | 0.0469 | -7.0464 | 0.0000 | *** |
| ***S. lycopersicoides*** | LA2776 | -0.1336 | 0.0311 | -4.2920 | 0.0006 | *** |
| ***S. lycopersicoides*** | LA4123 | -0.1313 | 0.0450 | -2.9176 | 0.1119 | ns |
| ***S. lycopersicoides*** | LA2772 | 0.1880 | 0.0298 | 6.3131 | 0.0000 | *** |
| ***S. lycopersicoides*** | LA1964 | 0.4522 | 0.0423 | 10.6881 | 0.0000 | *** |
| ***S. lycopersicoides*** | LA2951 | 0.5182 | 0.0463 | 11.2031 | 0.0000 | *** |
| ***S. pennellii*** | LA1303 | -0.2561 | 0.0324 | -7.9098 | 0.0000 | *** |
| ***S. pennellii*** | LA2657 | -0.1336 | 0.0229 | -5.8337 | 0.0000 | *** |
| ***S. pennellii*** | LA1656 | -0.1202 | 0.0396 | -3.0352 | 0.0779 | . |
| ***S. pennellii*** | LA1282 | -0.0897 | 0.0230 | -3.8936 | 0.0033 | ** |
| ***S. pennellii*** | LA1809 | -0.0774 | 0.0267 | -2.8946 | 0.1202 | ns |
| ***S. pennellii*** | LA2719 | -0.0714 | 0.0271 | -2.6405 | 0.2405 | ns |
| ***S. pennellii*** | LA0716 | -0.0397 | 0.0288 | -1.3792 | 0.9967 | ns |
| ***S. pennellii*** | LA1941 | 0.2596 | 0.0501 | 5.1787 | 0.0000 | *** |
| ***S. pennellii*** | LA2963 | 0.5285 | 0.0445 | 11.8656 | 0.0000 | *** |
| ***S. pimpinellifolium*** | LA2983 | -0.2359 | 0.0571 | -4.1295 | 0.0013 | ** |
| ***S. pimpinellifolium*** | LA4713 | -0.1162 | 0.0878 | -1.3232 | 0.9983 | ns |
| ***S. pimpinellifolium*** | LA1374 | -0.0457 | 0.0302 | -1.5143 | 0.9871 | ns |
| ***S. pimpinellifolium*** | LA2853 | -0.0209 | 0.0555 | -0.3770 | 1.0000 | ns |
| ***S. pimpinellifolium*** | LA2347 | -0.0174 | 0.0356 | -0.4882 | 1.0000 | ns |
| ***S. pimpinellifolium*** | LA1659 | -0.0172 | 0.0294 | -0.5839 | 1.0000 | ns |
| ***S. pimpinellifolium*** | LA1593 | 0.0622 | 0.0307 | 2.0227 | 0.7566 | ns |
| ***S. pimpinellifolium*** | LA1332 | 0.0895 | 0.0302 | 2.9637 | 0.0972 | . |
| ***S. pimpinellifolium*** | LA1348 | 0.1126 | 0.0319 | 3.5288 | 0.0142 | * |
| ***S. pimpinellifolium*** | LA1261 | 0.1890 | 0.0741 | 2.5516 | 0.2996 | ns |

Suppl. Table 5 *Solanum* accessions used in this study

| Accession | Solanum species |
|---|---|
| **LA0716** | *pennellii* |
| **LA1261** | *pimpinellifolium* |
| **LA1282** | *pennellii* |
| **LA1303** | *pennellii* |
| **LA1332** | *pimpinellifolium* |
| **LA1348** | *pimpinellifolium* |
| **LA1374** | *pimpinellifolium* |
| **LA1559** | *habrochaites* |
| **LA1593** | *pimpinellifolium* |
| **LA1656** | *pennellii* |
| **LA1659** | *pimpinellifolium* |
| **LA1721** | *habrochaites* |
| **LA1731** | *habrochaites* |
| **LA1753** | *habrochaites* |
| **LA1809** | *pennellii* |
| **LA1941** | *pennellii* |
| **LA1964** | *lycopersicoides* |
| **LA2128** | *habrochaites* |
| **LA2167** | *habrochaites* |
| **LA2347** | *pimpinellifolium* |
| **LA2409** | *habrochaites* |
| **LA2657** | *pennellii* |
| **LA2719** | *pennellii* |
| **LA2772** | *lycopersicoides* |
| **LA2776** | *lycopersicoides* |
| **LA2777** | *lycopersicoides* |
| **LA2853** | *pimpinellifolium* |
| **LA2864** | *habrochaites* |
| **LA2951** | *lycopersicoides* |
| **LA2963** | *pennellii* |
| **LA2983** | *pimpinellifolium* |
| **LA4123** | *lycopersicoides* |
| **LA4130** | *lycopersicoides* |
| **LA4713** | *pimpinellifolium* |
| **C32** | *lycopersicum* |

### Figures

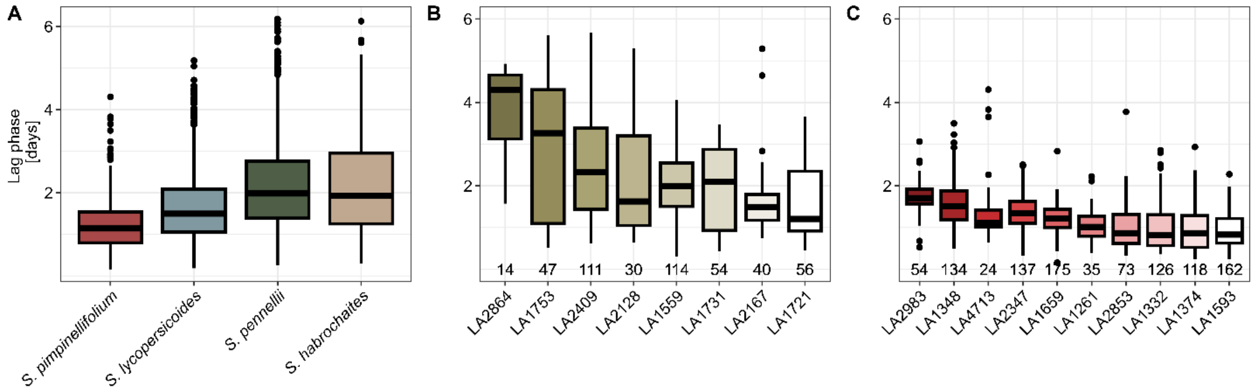


Suppl. fig. 1: **Lag-phase duration show different levels of variation depending on the host species A)** The lag phase duration (in days after infection) of *S. sclerotiorum* infection on *S. pennellii, S. lycopersicoides, S. pimpinellifolium* and *S. habrochaites* accessions. (B)Lag-phase duration of *S. habrochaites* accessions and C) *S. pimpinellifolium* accessions. The number on the x-axis indicates the count of individual leaves tested.

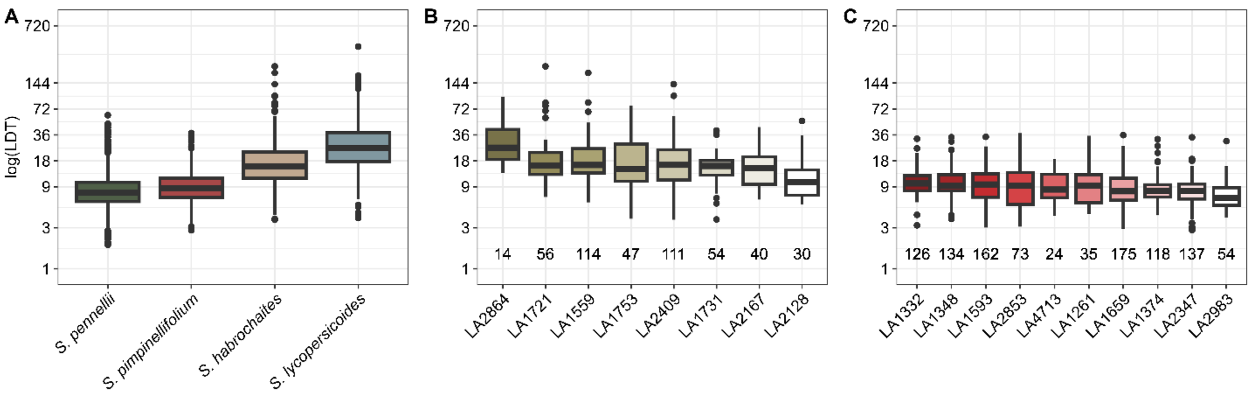


Suppl. fig. 2: **LDT show different levels of variation depending on the host species A)** The LDT (in hours) of *S. sclerotiorum* infection on *S. pennellii, S. lycopersicoides, S. pimpinellifolium* and *S. habrochaites* accessions. (B) LDT of *S. habrochaites* accessions and C) *S. pimpinellifolium* accessions. The number on the x-axis indicates the count of individual leaves tested.

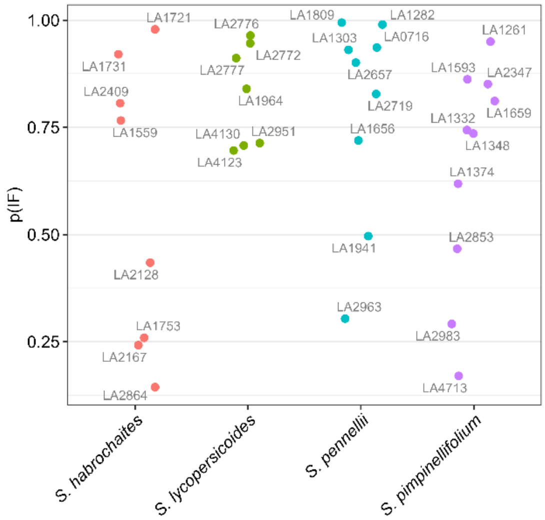


Suppl. fig. 3: **Per-accession infection frequency estimates. Values derived from a glm on three independent repetitions.**

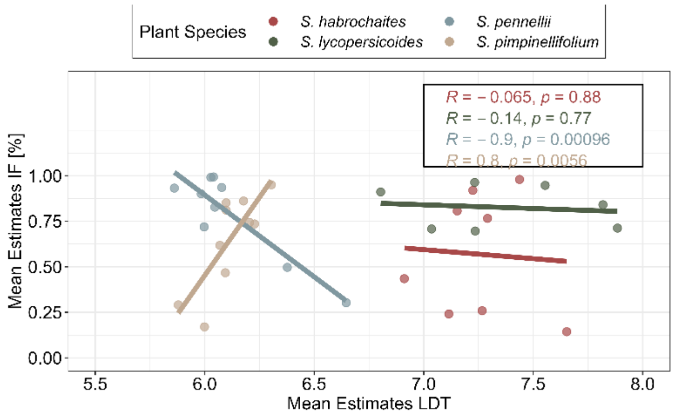


Suppl. fig. 4: **Correlation analysis between Infection frequency estimates and LDT.**

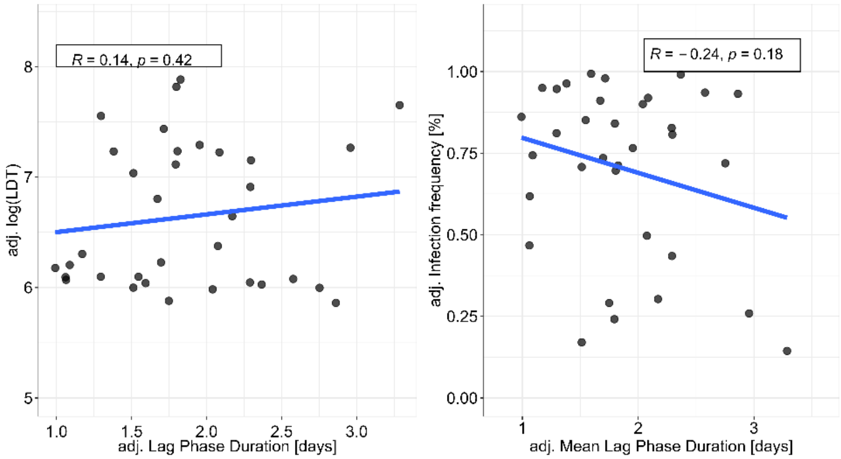


Suppl. fig. 5: **Pooled correlation analysis of all accessions.**

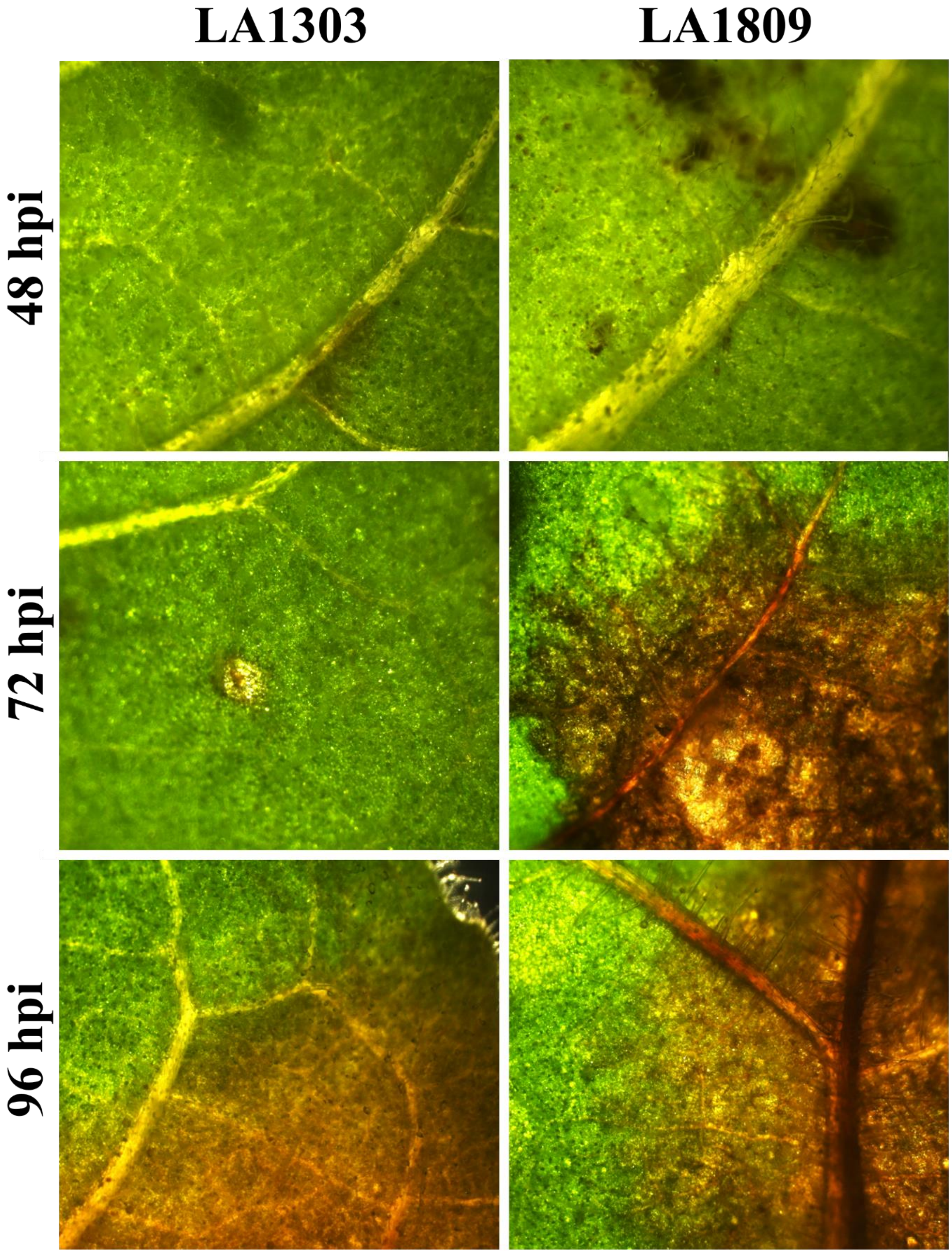


Suppl. fig. 6: **Bright light microscopy images of *S. sclerotiorum* infections on two *S. pennellii* accessions with different lag phase durations.**

Suppl. fig. 7: **Residual plot of lag-phase duration values.**

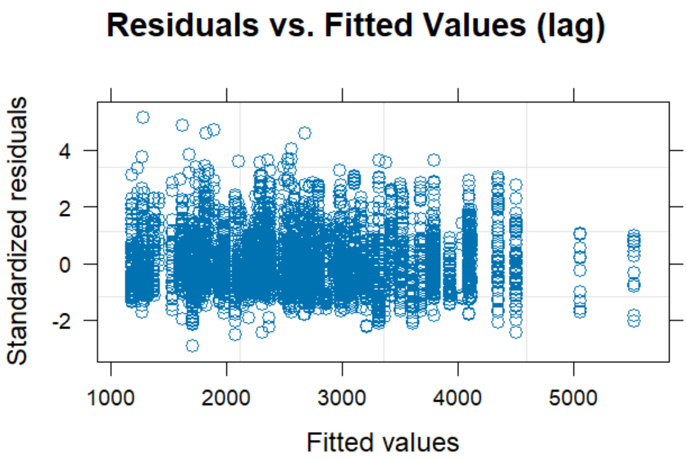


Suppl. fig. 8: **Residual plot of LDT values.**

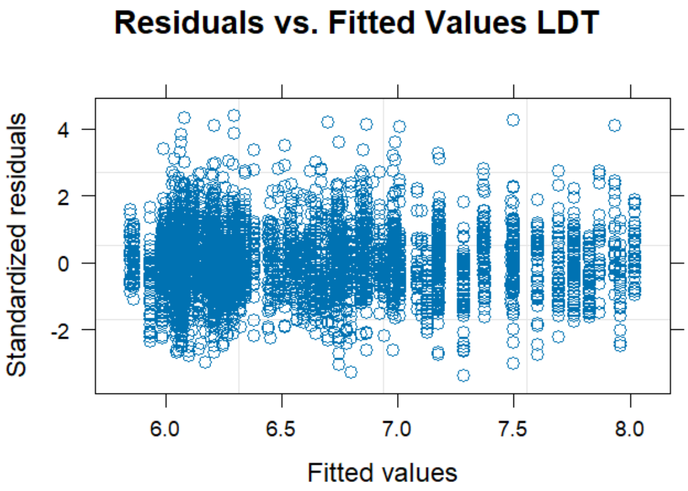
